# Disentangling the eco-evolutionary drivers of host-associated microbiota at the scale of an arthropod meta-community

**DOI:** 10.64898/2026.11.24.768847

**Authors:** Emma Fromm, Julien Cote, Léa Beaumelle, Elvire Bestion, Lucie Di Gesu, Uxue Suescun, Elodie Darnet, Lucie Zinger

## Abstract

While arthropod-associated microbiotas are known to be shaped by ecological and evolutionary processes linked to host traits and environmental context, their relative role at the scale of whole communities remains unclear. Here, we address this knowledge gap in a six-year experiment in semi-natural mesocosms, where both climatic variables and food web structure – via the presence of an apex predator – were manipulated. We tested the respective effects of host phylogeny, its trophic position, its mobility, as well as host climatic and biotic context on the diversity and composition of whole-body microbiota. Overall, 1145 composite samples in total encompassing 34 arthropod families, were characterized through DNA metabarcoding. We found a strong phylosymbiosis pattern, partly driven by endosymbionts and phylogenetically conserved host traits. However, we also found microbiota diversity to also respond to the climatic treatment and host ecological context – in particular prey and resource availability, albeit with a strength varying across hosts. For example, detritivores exhibited particularly high sensitivity to warming and interactive effects between the climatic treatment and resources. Overall, our findings highlight that the arthropod-associated microbiota is primarily structured by deeply conserved host traits or host-microbial associations shaped over long-term evolutionary time-scale, and to a lesser extent by ecological drivers such as trophic interactions or climate variability, operating over shorter timescales. Integrating these two temporal scales will be essential to accurately predict climate impacts on host-associated microbiota and to understand how these microbial shifts feedback on host physiology and ecosystem functions.

## Introduction

Arthropods are well known to be colonized by a variety of microorganisms (bacteria, virus, and fungi) on their body surfaces and within various organs and specialized cells [1]. These associations range from transient microbes to close and long-term associations [1,2], the latter being major drivers of arthropod metabolic physiological, and behavioural functions [3]. Microbial communities are assembled through simultaneous and partly confounded processes, including host filtering, environmental acquisition, dispersal and stochasticity [4,5], and disentangling these processes is key to predicting the contribution of microbiota to host function and their response to environmental change. At the long-term end of this continuum are primary obligate endosymbionts, which are often found within specialized cells (bacteriocytes), typically vertically transmitted, highly host-specific, and synthesizing nutrients lacking in the host’s diet [6], such as *Buchnera* in aphids and *Sulcia* in sap-feeding insects [7,8]. Arthropods may also harbor facultative endosymbionts that, while not essential for survival, can enhance host fitness, as suggested for Rickettsia strains in the whitefly *Bemisia tabaci* [9]. More broadly, arthropod-associated microbial communities, whether acquired vertically or horizontally, can significantly influence their host functions, adaptation, and resistance. For example, in attine ants, the host promotes competitive symbionts able to produce antibacterial molecules by providing high level of resources, thereby limiting the cuticle colonization by external pathogens [10].

As for any host-associated microbial communities, those in arthropods are governed by community assembly rules involving basic ecological and evolutionary processes (i.e. selection, dispersal, drift, diversification) mediated by host traits and environmental context [4,11,12]. A few meta-analyses support host taxonomy and diet as key and consistent drivers of gut and whole-body microbiota [13–17]. Beyond taxonomy, host evolutionary history also shapes microbiota composition in certain clades, including insects, mammals, and sponges [18,19]. More specifically, closely related species tend to harbour more similar microbiota, a pattern known as phylosymbiosis, resulting from long-term host–symbiont co-evolution and the recruitment (or exclusion) of microbial taxa by phylogenetically conserved life-history (e.g. mobility) or trophic (e.g. trophic guild or diet) and phenotypic traits [18,20]. Vertically transmitted endosymbionts - typically Wolbachia, one of the most studied endosymbionts in arthropods - may further reinforce this phylogenetic signal through their close co-diversification with host lineages [21]. However, the strength of the phylosymbiosis pattern varies across taxa, taxonomic scale and ecological contexts. For instance, while the phylosymbiosis pattern is detectable at multiple taxonomic levels in insects (from species to order), it weakens at broader scales [15], and in some cases, is even very weak or absent [16,22]. In orthopterans, the microbiota composition is better explained by host species identity and habitat than by host phylogeny [23].

On shorter ecological time scales, abiotic and biotic conditions, including anthropogenic ones, further shape arthropod microbiota [16,17,24]. Among them, climate and trophic interactions are of particular interest, as the loss of predators and/or preys and climate warming are two cooccurring components of global change [25]. Among biotic interactions, prey availability can shape dietary intake and thus microbiota composition, as shown in predatory beetles who exhibited higher microbial richness when prey diversity increased [26]. Reciprocally, predators can influence their prey microbiota indirectly. First, predation risk can influence prey’s microbiota composition via stress-induced physiological changes, as observed in perch and mussels [27,28], and suggested for in arthropods [29]. Alteration of host density through predation can also affect microbial transmission, especially in social insects through increase or disruption of contact interactions such as mating and coprophagy [30,31]. Lastly, species’ movement abilities can also influence their microbiota, with flying terrestrial invertebrates accessing a wider range of microhabitats and microbial pools compared to less mobile insects that rely more on local microbes [32,33]. Among abiotic factors, temperature is well-known to influence host-symbiont relationships across various invertebrate clades [34]. Experimental warming has been shown to induce shifts in microbiota composition in *Drosophila* [35], the green stinkbug (*Chinavia hilaris*) [36] and in aphids [37]. In natural environments, temperature effects are likely multifactorial, combining direct influences on microbial communities with indirect effects through host physiology, resource availability and/or intake, predator–prey dynamics, and community composition [38].

Yet, the relative importance and interplay of all these factors remain poorly understood, largely due the incapacity of disentangling these often-confounding factors in field studies, and the difficulty to manipulate all of them experimentally under ecological realistic conditions. So far, studies have focused on specific model systems, primarily hemipteran species such as the aphid–*Buchnera* associations or the leafhoppers-*Sulcia* associations [39]. While such studies are starting to be extended to other arthropod taxa, including isopods [40,41], spiders [42], and Chrysomelidae [43], the combined complexity of the all the factors outlined above remains to be disentangled at the scale of an arthropods community.

Here we investigate the evolutionary and ecological factors shaping the microbiota of arthropods at the scale of a whole meta-community. We rely on a six year-long warming experiment in a semi-natural setup consisting of mesocosms where we manipulated climatic conditions, as well as biotic assemblages, through the introduction or not of an apex generalist predator, the common lizard (*Zootoca vivipara*) [44]. Previous work demonstrated that the presence of this predator alters the plant and invertebrate community [45]. We monitored arthropods’ abundance as well as a wide range of biotic and climatic variables to assess their link with arthropods’ microbiota, characterized through DNA metabarcoding. We then first provide a description of the microbiota composition associated with the 34 most occurrent arthropod families across mesocosms. Second, we quantify the effects of host evolutionary history and associated host traits relative to that of biotic/abiotic conditions on microbiota communities. We further examine how these local abiotic/biotic conditions influence the distribution of two major arthropod endosymbionts: *Wolbachia* and *Rickettsia*. Finally, within ubiquitous arthropod families, we assess the effect of various biotic and climatic factors on microbiota diversity and composition. We hypothesize that host phylogeny, host trophic guild and mobility will be major determinants of the microbiota diversity and composition.

## Material and Methods

### Experimental system

We conducted a six-year-long experiment (2015-2021) in the Metatron experimental setup (Moulis, Ariège, [44]). The Metatron consists of 10 × 10 m mesocosms bordered by buried tarpaulins (50 cm depth) and covered with insect-proof nets. Each mesocosm functions as a mini ecosystem supporting naturally assembled vegetation and arthropod communities. Here, we used 24 mesocosms in total to conduct our experiment [46].

### Climatic treatment and monitoring

Climatic conditions were manipulated using shutters that automatically close when ambient temperature exceeds either 28°C (present-day) or 38°C (warming). This manipulation is thus active only when ambient temperature becomes high, i.e. in summer, and has been shown to produce a realistic 1-3°C warming effect depending on daily, seasonal, and inter-annual climate fluctuations [44]. We monitored various variables throughout the experiments using sensors located in each cage. From these, we extracted minimum, mean and maximum temperatures for each season of each year in each cage. We also calculated the number of hours when temperatures exceeded 34°C and 38°C each year. We summarized these climatic variables with a principal component analysis (PCA). The first axis (explaining 33.5% of variance, Fig. S1) was used as a descriptor of the climatic treatment in the analyses. We used here climate from the preceding year (*t*-1) (hereafter Climate (*t-*1)) to explain microbiota diversity and composition at year *t* because the sampling was conducted at the beginning of summer, when the yearly climatic treatment only starts to be active.

### Lizards’ treatment and populations monitoring

An apex predator treatment was implemented by introducing lizard individuals into 16 mesocosms (8 per climatic modality), with 8 predator-free control mesocosms (4 per climatic modality). *Zootoca vivipara* is an opportunistic insectivorous predator that feeds on a diverse range of arthropods [47–49] present in the Metatron [45,50]. Its populations were monitored twice yearly (May and September) from 2015 to 2021 using standardized captures sessions [46,51], during which individuals were identified, weighted and aged. To analyse the effects of population size and age structure of apex predators, we calculated for each mesocosm and year: the lizard abundance (individuals per mesocosm), the total biomass (sum of individual masses), and the biomass to abundance ratio (total biomass divided by abundance). Because adults have higher biomass than juveniles, the biomass-to-abundance ratio reflects the average body size and age distribution.

### Plant communities monitoring

Plant communities were surveyed in June 2017, 2018, and 2021 along standardized linear transects (3 m x 1.5 m) within each mesocosm. All plant species present within each transect strip were identified and recorded. From this, we retrieved the total species richness in each mesocosm for each sampled year. To assess the consistency of plant community composition over time, we evaluated the repeatability of species richness across years within mesocosms. Given the high repeatability (R=0.63), we estimated the species richness for the missing survey year (2016, 2019 and 2020) by averaging the values of the same mesocosm from the two closest surveyed years, except for 2016 for which we only used data from 2017.

### Arthropods sampling and preparation

Arthropods were sampled twice a year: in June, when the climatic treatment starts to be active, and in September the same year after the treatment becomes inactive. In June, pitfall traps targeting ground-dwelling arthropods and sweep-netting targeting vegetation-associated taxa were deployed per mesocosm. Only sweep-netting was conducted in September. These sampling methods have been shown to provide adequate sampling coverage [50,52]. Specimens were preserved at –20 °C, identified to species or genus when possible (otherwise at family level) and sorted to family level for analyses. We chose this grain of description for arthropod hosts and their microbiota to ensure their presence across mesocosms and treatments.

To characterize arthropod community composition within each mesocosm, arthropod families were assigned to one of four trophic guilds (detritivorous, phytophagous, predator, or omnivorous) based on literature, with omnivorous groups excluded due to their low abundance and ambiguous isotopic signatures. Families whose members have mixed feeding strategies were assigned to the trophic guild corresponding to their predominant feeding strategy locally. For each mesocosm, we thus report (1) the total arthropod abundance per trophic guild, obtained by summing abundance values in September after the climatic treatment, and in June, at the beginning of the treatment the year after, in order to capture respectively short (≤ 4 months) to longer-term (≥ 8 months) effects of the climatic treatment, and (2) the Chao richness [53] per trophic guild. Data were combined over the two sampling periods (June at year t and September at year t-1) because we observed consistent responses to treatments when we initially modeled them separately, allowing for a simplified model structure in downstream analyses.

For isotopic and microbiota analyses, we selected 34 arthropod families (Table S1, Fig. 1) representing families in each trophic group ubiquitous across mesocosms, i.e. 2 detritivorous families (Julidae, Isopoda), phytophagous Cicadellidae, predator Lycosidae, and the most abundant families in each mesocosm. As most of these families are frequently found in lizard diets [47–49], we expect them to be influenced by the lizard biomass. For each family, mesocosm and year, individuals were pooled into composite samples depending on available samples and individuals biomass (sample size = 1-23 individuals, typically 3) to account for intra-family variability. Due to our sampling method, within a given mesocosm, several samples of the same family were available for comparison only for detritivores. For each sample, individuals’ bulks were cleaned with distilled water, dried with Kimtech paper, lyophilized for 7 days, and ground prior analysis. Sampling blanks were included for each processing batch during sample sorting and pooling to monitor potential contamination. We thus obtained a total of 1145 arthropod powders. Prior statistical analysis we ensured that the varying sample size we worked with did not influence bacterial diversity or phylogenetic composition metrics (Fig. S2).

**Figure 1.**
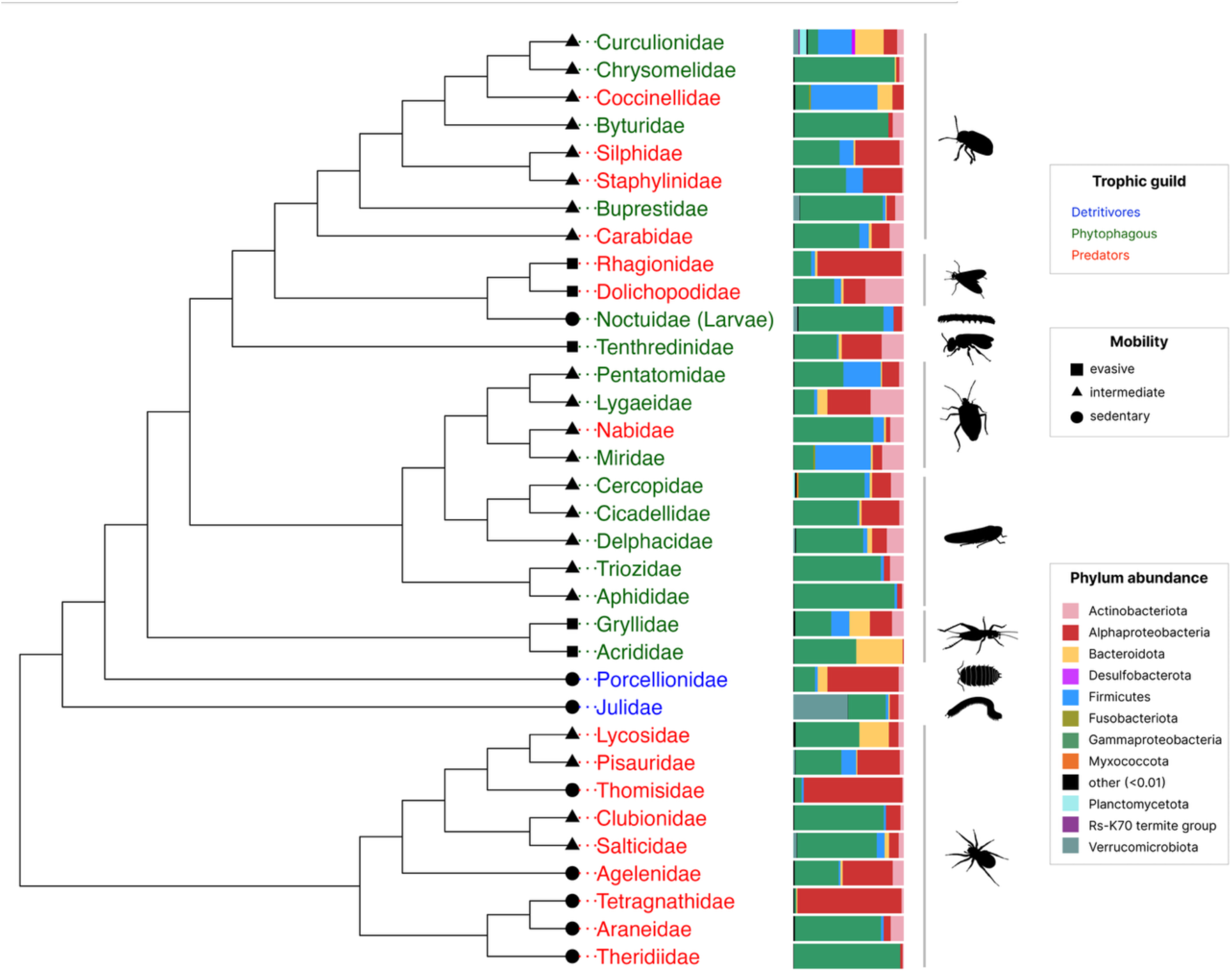
Overview of the studied arthropod hosts showing their phylogenetic relationships, their ecological traits, and their microbiota composition. Tips are coloured by trophic guild and shaped by mobility status. Relative abundances of dominant phyla are shown as bar diagrams, summed across hosts within each family.

### Arthropod phylogenetic and functional characteristics

To characterize the evolutionary relatedness of the 34 arthropods families selected, we retrieved a phylogenetic tree of all arthropods families based on TimeTree [54] using Treehouse [55] (Fig. 1). As the Liviidae family was absent from TimeTree and because our primary objective was to estimate the broad relatedness between host families rather than to study their precise phylogenetic relationships, the missing family Liviidae was artificially substituted with its sister group, Triozidae. From this subtree, we generated a pairwise phylogenetic distance matrix between families using the ape package.

To identify the trophic position of arthropod families, lyophilized arthropod powders were split into two subsamples, one used for microbial analyses (see below) and the other for δ¹³C and δ¹⁵N measurements, which were conducted by the Cornell Stable Isotope Laboratory (Ithaca, USA). Carbon and nitrogen isotope values were normalized to the Vienna Pee Dee Belemnite (VPDB) and atmospheric air standards, respectively [56]. Host mobility was classified at the family level as sedentary, intermediate, or evasive following [57], refined using [58,59], and validated using wing morphology from specimen photographs and literature. For Araneae, for which wing morphology cannot be used to assess mobility, classification was instead based on their predominant locomotory strategy: web-building families were considered sedentary, whereas active-hunting taxa were considered intermediate.

### Microbiota sampling and molecular analysis

We characterized the whole-body microbiota of the 34 arthropods families from the other subsample of the arthropod powders. Total DNA was extracted with the Qiagen Dneasy Blood & Tissus Kit Tube (Qiagen GmbH, Hilden, Germany). We conducted PCR amplification using the BACTB-F (50-GGATTAGATACCCTGGTAGT-30) and BACTB-R (50-CACGACACGAGCTGACG-30) primers [60], which target a ∼250-bp-long marker located in the v5–6 region of the bacterial 16S rRNA gene. Both primers were labelled at their 5’ ends with two different 8 nt tags to discriminate PCR reactions.

PCR reactions were conducted in triplicate for each sample with AmpliTaq Gold DNA Polymerase in 20 μl reaction volumes (See Supplementary Appendix A for detailed protocols). The PCR products, along amplified sampling, extraction and PCR blank controls were then pooled and purified with the QIAquick PCR purification Kit (Qiagen GmbH, Hilden, Germany) to prepare nine libraries using Illumina TrueSeq protocol without the enrichment step to reduce the amount of tag-jumps. The libraries were sequenced on a NovaSeq6000 platform using 2×250 bp paired-end chemistry at the GetPlage platform (Toulouse, France).

### Bioinformatic analysis

Sequence analysis was conducted using OBITools v1.2.11 [61,62], the OBITools v4 (https://github.com/metabarcoding/obitools4), and Sumaclust [63] softwares in a Snakemake [64] workflow. Paired-end reads were aligned, assigned to their respective samples, dereplicated, and filtered when exhibiting low alignment scores (<40), a length < 100 bp, ambiguous nucleotides, errors on tags, more than two mismatches on primer sequences, and when they were singletons. We further denoised the dataset using the *obiclean* (v4) program, a graph-based approach detecting PCR errors [61], using default parameters. Because several reports suggest that this level of resolution tend to overestimate the diversity in metabarcoding data, including with 16S markers due to remaining errors or intra-genomic variability [65–68], the denoised sequences were clustered into Operational Taxonomic Unit (OTU) using Sumaclust with a 97% similarity threshold. The most abundant sequence in each OTU was designated as its representative, with its abundance reflecting the sum of all member sequences. To assign each OTU a taxon, we used the SILVA database (release 138.2) and the taxonomic assignment tool from the SILVAngs pipeline [69]. Taxonomic assignments having a probability < 80% were considered as unreliable. Finally, we inspected the dataset quality and curated data from sampling, extraction, or PCR contaminants and potential tag-jump based on experimental blank control using the metabaR package [70]. Sequences assigned to chloroplasts and mitochondria were also excluded.

Our final dataset comprised 1 329 446 OTUs and 140 732 744 reads corresponding to 1145 samples including 438 detritivores, 359 phytophagous and 348 predators. We covered 34 dominant arthropods families in the Metatron, including 2 detritivorous, 16 predatory and 16 phytophagous families (Table S1, Fig. 1).

### Diversity measurement

Rarefaction curves (Fig. S3) generated with the *vegan* package [71] indicated an adequate sequencing coverage, especially when quantifying the diversity with the exponential of the Shannon index (exp(H)). As this index has been shown to reduce biases associated with insufficient sampling [72] and to reduce the influence of rare molecular artifacts that may remain in the dataset [73], we used it for downstream diversity analyses.

### Phylogenetic placement

Preliminary analyses showed no overlap in microbial taxonomic composition across host families, which limits the interpretation that can be drawn from this diversity facet. Because the tested drivers of arthropod microbiota are also likely to leave evolutionary imprints at both small and deep time scales, we analysed instead their diversity through a phylogenetic prism. We used a phylogenetic placement approach for short 16S rRNA sequences rather than de novo reconstruction, which is known to be less reliable [74]. Briefly, OTUs were aligned and placed onto a reference phylogeny of 20,000 bacterial and archaeal 16S rRNA genes using EPA-ng within the first two steps of the PICRUSt2 workflow [75,76]. Placement quality was assessed with Gappa [77], and Likelihood Weight Ratio (LWR) values indicated high-confidence placements for most OTUs (Fig. S4). Pairwise microbial phylogenetic dissimilarities between hosts was calculated using the Kantorovich–Rubinstein (KR) distance.

### Statistical analysis

#### Phylosymbiosis pattern analyses

To explore patterns of phylosymbiosis, we measured the correlation between host phylogenetic distances and bacterial phylogenetic dissimilarities (KR distance) using a Mantel test (999 permutations, Pearson coefficient) [79]. We then partitioned the variance in bacterial KR distances between host phylogeny, phylogenetically conserved host trait (trophic isotopic position and host mobility) [32,80] and mesocosm identity using a hierarchical variance partitioning model (*rdacca.hp*, [81]), allowing to tease apart evolutionary drivers from ecological drivers in the assembly of arthropod microbiota.

Since the model cannot directly handle distances, we performed an Edge Principal Component Analysis (Edge PCA) with Gappa on the bacterial distances between samples, based on the KR index [82]. Edge PCA is specifically designed for phylogenetic placement data and identifies which reference tree taxa contribute most to compositional differences among samples. We then extracted the projections of the samples onto the principal components as the response variable in the model. Similarly, host phylogenetic distances were analysed via a Principal Coordinates Analysis (PCoA), which provides an ordination of sample relationships based on those distances. We evaluated the trophic isotopic position using the isotopic signature dissimilarities between arthropod families by conducting a PCoA on Euclidean distance of the mean values of δ¹⁵N and δ¹³C between each family (Fig. S5). For all ordinations, we retained the axes cumulatively explaining 90% of the total variation, corresponding to 11 axes for the bacterial Edge PCA and for host phylogenetic PCoA, and 2 axes for trophic isotopic position. For host mobility and local habitat (i.e. mesocosm/year), we created dummy variables, with each level represented by a column, minus one reference level (sedentary and one mesocosm from one year), which is implicitly encoded by the sum of the other variables.

#### Contribution of endosymbionts to the phylosymbiosis pattern

We further investigated more specifically the distribution of two widespread arthropod endosymbiont genera (Table S1): *Wolbachia* and *Rickettsia* [83]. To account for the host phylogenetic effects and co-varying host traits, we included as fixed effects: the host phylogenetic distinctiveness (i.e. host position along the first host phylogeny PCoA axis), the host mobility and trophic isotopic position. As these bacterial genera can be transmitted both vertically and horizontally [84,85], we examined whether their presence at year *t* was influenced by the potential host carriers and dispersal vectors by including global arthropod richness and lizard abundance. Finally, we include the effect of climate conditions at year *t-1* as it could modify the interaction between a host and its endosymbionts [36,86,87]. Random effects include host family, year of sampling and mesocosm identity. When only one single read was assigned to *Wolbachia* or *Rickettsia* in a sample, the genus was considered as absent. All candidate generalized linear mixed models were ranked by AIC, and models with ΔAIC < 2 were averaged (MuMIn; [88]). We extracted conditional estimates, standard errors, z-values, relative importance (RI), and p-values. Model assumptions were validated graphically by assessing residual normality and homoscedasticity.

#### Drivers of arthropod microbiota taxonomic diversity

As host phylogeny strongly structured microbiota composition, we investigated how climate and environmental factors shape microbiota variation within host taxa at shorter time scale. We focused on microbiota diversity for ubiquitous families with sufficient sample size (Cicadellidae, Isopoda, Julidae, and Lycosidae). As a complementary analysis, we extended this approach at the trophic group level to test whether the patterns found within families were conserved (Supplementary Appendix B). For each family, we fitted a linear mixed model using a model-averaging approach. All variables were scaled [89] to estimate the overall effects over the dataset. Full linear mixed models (lme4, [90]) included as fixed effects (i) the abundance and richness of each arthropod trophic guild (i.e. conspecific abundance and/or predator and prey abundance and richness), (ii) plant species richness (i.e. plant resource diversity), and (iii) lizard abundance and biomass (i.e. predation pressure). These variables were modelled in interaction with the climatic treatment Climate (t-1, i.e. year before sampling) to test for climate-dependent effects. Because the effect of climate on microbiota diversity could be nonlinear [91,92] and we added a quadratic term (Climate (t-1)²) as fixed effects to account for this effect. Random effects included year, mesocosm identity and were removed when not explaining any variance from the final models. Microbiota diversity was analysed with the same averaging procedure than for endosymbionts analyses but using a linear mixed model.

#### Drivers of arthropod microbiota phylogenetic beta diversity

We used Generalized Dissimilarity Models (GDM, [93,94] to estimate the variance in microbiota communities phylogenetic dissimilarity, based on the normalized KR index explained by the same variables as described in the model for diversity for ubiquitous families and trophic guilds.

## Results

### Microbiota taxonomic composition of a whole arthropod metacommunity

At the phylum level (Fig. 1), Gammaproteobacteria were the most abundant clade in many arthropod families of isopods (Isopoda), hoopers (Cicadellidae, Cercopidae, Delphacidae), beetles (Curculionidae, Staphylinidae, Coccinellidae), spiders (Lycosidae, Araneidae, Pisauridae, Salticidae, and Tetragnathidae), bugs (Nabidae), and flies (Dolichopodidae), consistent with different reports [40,42,95,96]. Alphaproteobacteria were the most prevalent in Carabidae and Chrysomelidae (beetles) as observed previously [13,43], and in the Tenthredinidae family (flies), while Firmicutes dominated in Julidae (millipedes) and Gryllidae (crickets). The microbiota composition at the genera level for the most abundant host families (Table S1) was also consistent with previous reports, with the dominance of *Sulcia,* the presence of *Pectobacterium* in Cicadellidae [8] and the high abundance of *Sodalis* in Cercopidae [97]. Well-known endosymbionts were also highly prevalent and abundant across multiple arthropods families, with *Wolbachia* in Dolichopodidae, Coccinellidae, Nabidae, Staphylinidae, Chrysomelidae, Gryllidae, and Delphacidae while Rickettsia dominated in Pisauridae, Lycosidae, Chrysomelidae, Curculionidae and Delphacidae. Finally, *Rickettsiella* was found in almost all host families except in Salticidae,

### Phylosymbiosis and its underlying drivers

We found a significant correlation between host family phylogenetic distance and bacterial phylogenetic compositional dissimilarities (Fig. 2A), hence revealing the existence of a phylosymbiosis pattern. We further show that the bacterial phylogenetic dissimilarity pattern is primarily driven by host evolutionary history. Host trophic position and mobility counted for a smaller, yet noticeable amount of variance, much of which overlaps with host phylogeny, due to the phylogenetic conservatism of these traits. This suggests that part of the phylosymbiosis pattern results from conserved host ecological features. Finally, the pure effect of local habitat was small, whereas its shared contribution with host phylogeny accounted for a non-negligible proportion of the variance (Fig. 2B, 2C). This shared fraction of variance likely reflects a non-homogeneous distribution of host clades across mesocosms – potentially driven by the climatic and biotic treatment. Such a filtering would indicate that local environmental effects are partly modulated by host phylogenetic constraints. Overall, our model explained 86% of the variance, indicating that the variables considered here captured a substantial proportion of the variation in bacterial phylogenetic composition.

**Figure 2.**
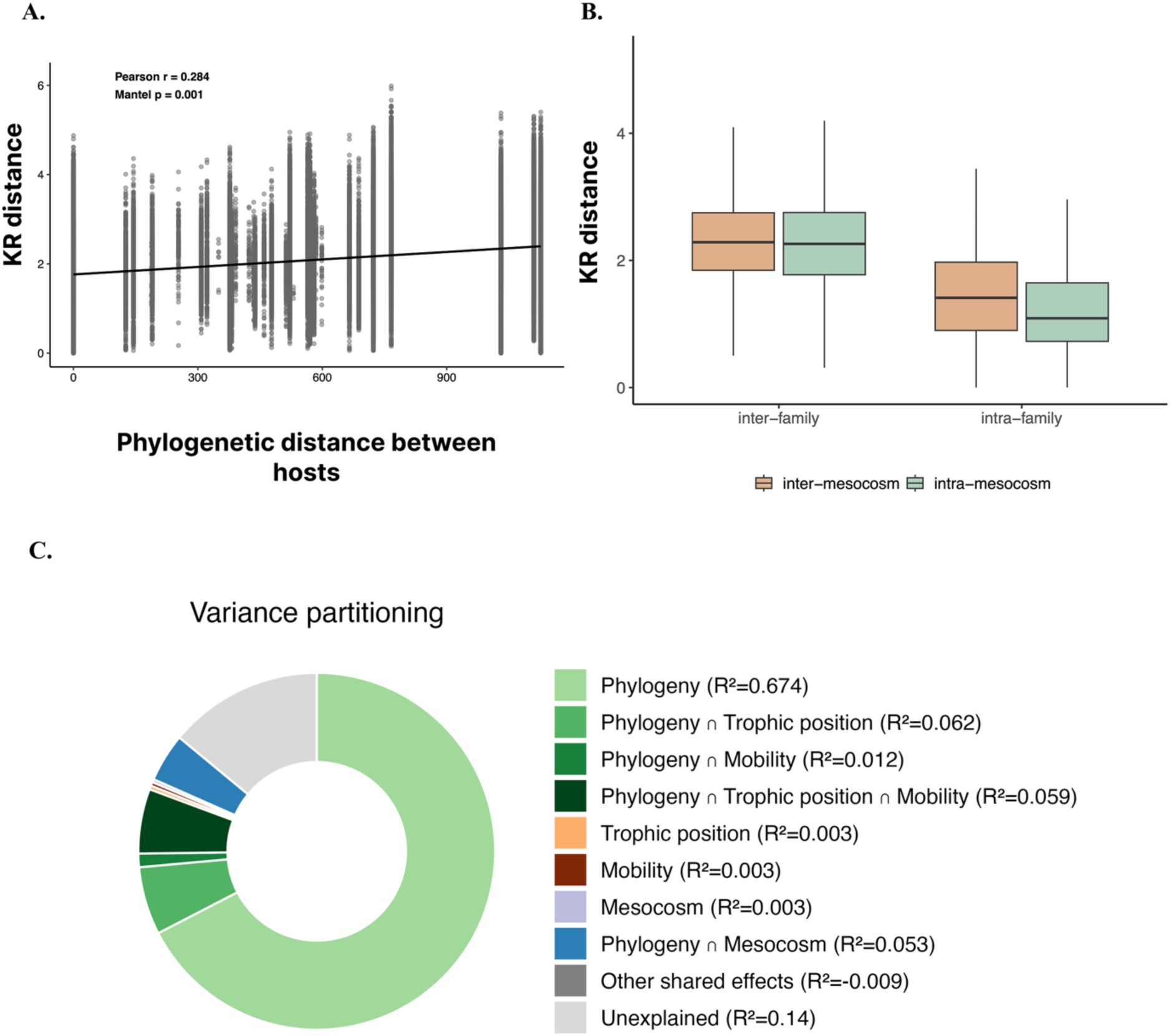
Bacterial community phylogenetic dissimilarity patterns at the scale of the arthropod metacommunity. **A.** Phylosymbiosis pattern, i.e. correlation between host phylogenetic distances and bacterial community phylogenetic distances. The Mantel test (999 permutations) was used to calculate the correlation coefficient (Pearson r) and associated p-value. **B.** Within-year phylogenetic dissimilarities of microbiota communities across all sampled arthropod families, comparing samples from the same or different host families and mesocosms. **C**. Contribution of host phylogeny, isotopic trophic position, mobility and mesocosm (mesocosm by year) and their shared effects in explaining the variance of bacterial phylogenetic distances, obtained with a hierarchical variation partitioning analysis.

### A focus on two known endosymbiotic genera

*Wolbachia* and *Rickettsia* were detected in a large proportion of samples (Table S1), with *Wolbachia* present in 991 out of 1145 samples and *Rickettsia* in 822 out of 1145. The probability of *Wolbachia* presence was positively associated with the PCoA1 of host phylogenetic distance (Table S2). Warming and host mobility were retained in the best averaged model, respectively with a high and low RI, even though p-value were not significant (Table S2). In contrast, *Rickettsia* presence was unaffected by host phylogeny but was negatively influenced by climatic conditions and positively by arthropod richness (Table S2).

### Unravelling host microbiota ecological drivers

In isopods (*Isopodae*), we found positive interactions between climate and detritivore abundance, as well as between climate and phytophagous diversity (Table S3), with a positive climatic effect on microbiota diversity restricted to habitats with high detritivore abundance (Fig. 3D) or phytophagous richness (Fig. 3C). This effect was cancelled in habitats with low detritivore abundance and even reversed in habitats with low phytophagous richness. Julidae’s microbiota diversity was reduced by warmer climates, and we found a negative interaction between climate and δ¹⁵N with stronger negative warming effects for individuals with higher trophic position (Fig. 3A, 3B, Table S3). In leafhoppers (*Cicadellidae*), we found a positive nearly significant quadratic effect of climate on the microbiota diversity (Fig. 3A, Table S3). Lycosidae microbiota did not respond to the climatic treatment but was negatively influenced by δ¹⁵N values (Table S3).

**Figure 3.**
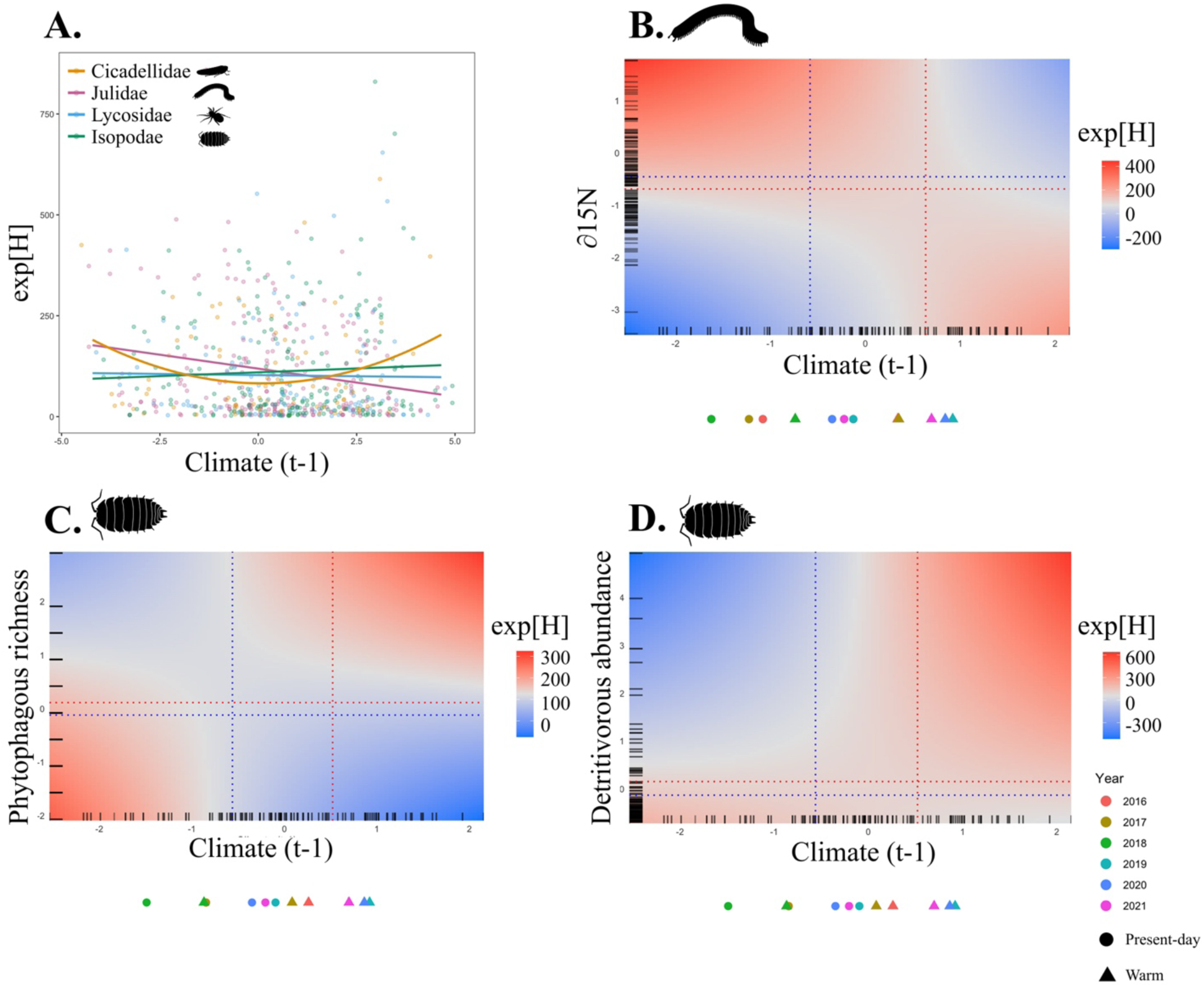
Ecological drivers of arthropod microbiota diversity for ubiquitous host families representatives of different trophic guilds. **A**. Overview of the response of microbiota diversity (exp [H]) to climatic conditions for each host family. **B**. Predicted response of Julidae (detritivorous host family) microbiota diversity to Climate (t-1) and ∂^15^N. **C.** Predicted response of Isopodae (detritivorous host family) microbiota diversity to Climate (t-1) and phytophagous richness. **D**. Predicted response of Isopodae microbiota diversity to of Climate (t-1) and detritivorous abundance. Colour gradients indicate predicted values of taxonomic diversity (exp[H]), and rugs represent observed values along both axes. Global mean and yearly means are shown for both present-day (dotted blue lines) and warm climate (dotted red lines) treatments. The corresponding statistical analyses are provided in Table S2.

The GDM models explained only a small fraction of the total deviance in bacteria phylogenetic dissimilarity patterns (ranging from 6% to 9%). In isopods (*Isopodae*), bacterial phylogenetic compositional changes were most strongly influenced by predator abundance, lizard biomass-abundance ratio and year of sampling. In millipedes (*Julidae*), the predator richness, abundance of detritivores and climate emerged as the primary drivers, whereas in wolf spiders (*Lycosidae*) the two best predictors were again predator abundance and phytophagous richness. Finally, for leafhoppers (*Cicadellidae*), the climate dominated, with the abundance of detritivores also contributing substantially (Fig. 4).

**Figure 4.**
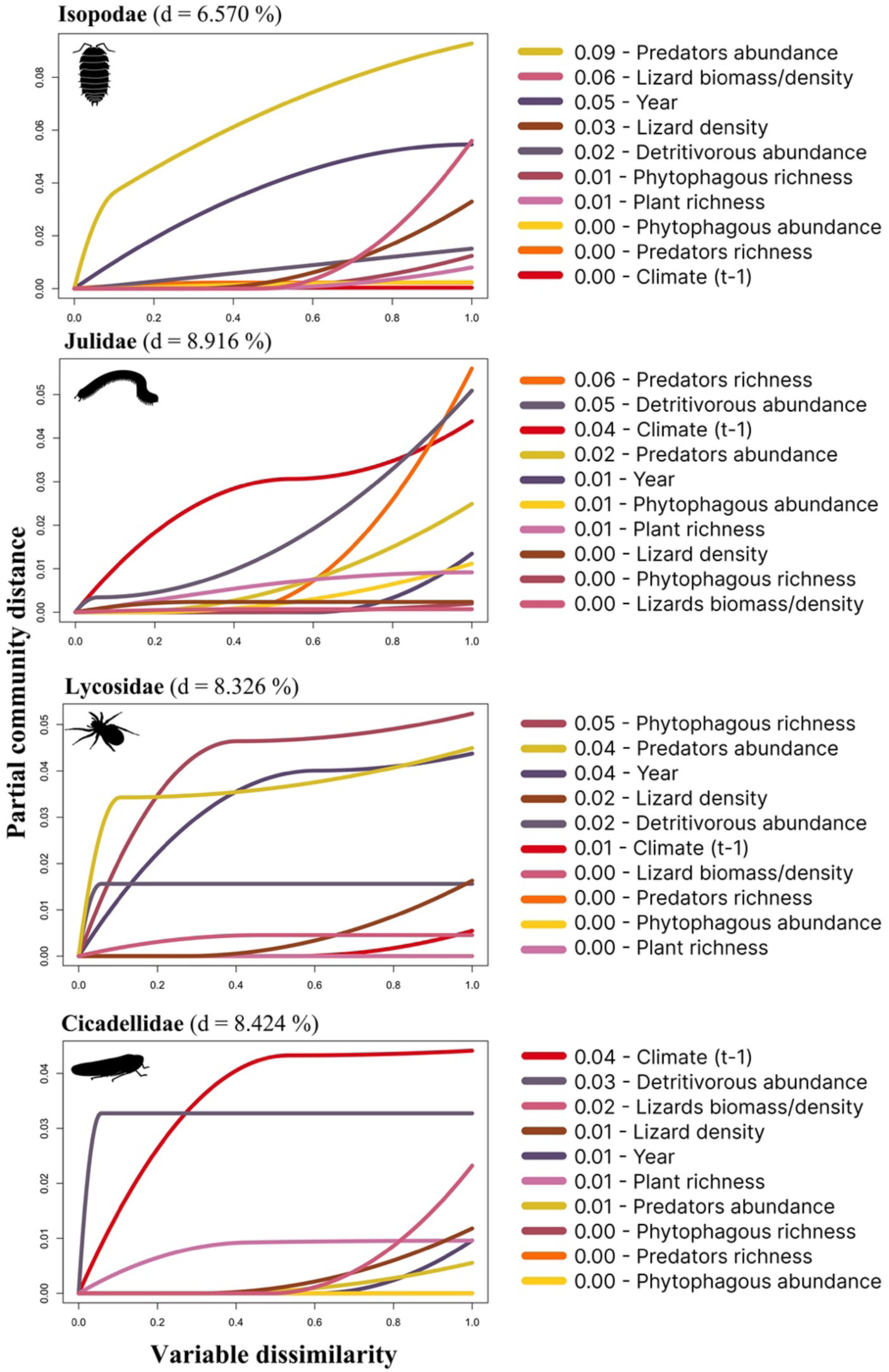
Drivers of Arthropoda microbiota phylogenetic dissimilarities as identified with Generalized dissimilarity models (GDMs). The GDMs splines for each ubiquitous arthropod families: Isopodae (detritivorous), Julidae (detritivorous), Lycosidae (predators) and Cicadellidae (phytophagous) are represented. The deviance (d) explained by the model is indicated along the host family. We used Climate (t-1), arthropods richness and abundance variables, as well as lizards abundance and plant richness as predictors. The shape of the spline shows the rate of community composition changes (y-axis) as the predictor variables (x-axis) increases. Predictors variables are scaled.

## Discussion

Here, we address host microbiota assembly processes at an unprecedented scale; encompassing an entire arthropod metacommunity maintained under semi-natural conditions. The experimental approach and close monitoring of climatic and biotic conditions provides a unique opportunity to disentangle how Arthropoda microbiota is shaped by host phylogeny, climate-driven environmental change and host biotic context (i.e. prey resources and predator pressure).

Bacterial communities strongly varied across arthropod families, consistent with the literature [42,95,98]. We further identified a clear phylosymbiosis pattern. While phylosymbiosis has been observed across diverse animals including insects, mammals, birds, fishes [18,19], the strength of this pattern seem highly variable depending on the breadth and identity of the taxa considered, as well as the ecological context [15,16,23]. Here, we observed this pattern is remarkably strong at the scale of 34 arthropod families of a single meta-community, despite considerable variability of the ecological context (biotic and abiotic) over the six years of the experiment. Such a robust pattern can be explained by our whole-body microbiota approach. Unlike many other studies, a whole-body approach increases the chances of capturing signal form vertically transmitted endosymbionts, which are more stable and linked to their hosts compared to the gut microbiota, whose composition can vary depending on the prey intake/identity [23].

This pattern suggests that the phylosymbiosis signal observed here arise from microbiota co-evolution with the host. Accordingly, we found a strong pure contribution of host phylogeny to the variance of bacterial phylogenetic dissimilarities, as well as higher probability of *Wolbachia* presence in phylogenetically close hosts. *Wolbachia* being a genus notoriously co-evolving with a range of arthropod hosts [99,100], it likely contributed to the phylosymbiosis signal observed here. In addition, such a signal can result from ecological filtering caused by phylogenetically conserved host traits, that would recruit similar microbial taxa from the environment [79]. Among host traits, host trophic mode/diet guild and dispersal abilities frequently show a strong phylogenetic signal [32,101] and have been repeatedly suggested to be involved in the assembly of animal host microbiota. Our analysis confirms that host isotopic trophic position and mobility covary with host phylogeny and contribute to the observed phylosymbiosis pattern, even though these confounding effects were less pronounced than the pure phylogenetic one. It is likely that other unmeasured phylogenetically conserved traits act as ecological filters, thereby contributing to the substantial pure phylogenetic effect.

When considering microbiota diversity instead of their phylogenetic dissimilarities, no correlation with host phylogeny or trophic position was found, suggesting that the phylosymbiosis signal observed here is more due to a phylogenetic turnover rather than a change in diversity [102]. The lack of differences in microbiota diversity among host isotopic trophic position and functional groups contrasts with previous studies reporting higher diversity in omnivores and detritivores than in herbivores and predators [14,17,101]. This discrepancy may again be primarily explained by our whole-body approach rather than a gut one, as the gut is more strongly influenced by ingested material. Indeed, our results are consistent with a previous study across various insects’ species showing that diet is a poor predictor of whole-body microbiota [15]. Similarly, we did not detect a clear effect of host mobility on bacterial community composition. Previous evidence of host mobility-driven microbial turnover has been observed during long-distance dispersal events between species [103] or at the intraspecific level in relation to individual locomotor activity [104]. Our analysis focused on differences among insect families and mobility effects may operate at finer taxonomic or behavioural scales than those captured here.

Finally, we also detected a non-negligible shared effects of local habitat (i.e. mesocosm identity) and host phylogeny in the phylogenetic composition of arthropods microbiota. This shared effect suggests that host lineage may modulate how microbial communities respond to local environments. In line with this, we observed a lower presence probability of *Rickettsia* under warmer conditions. While *Rickettsia* symbionts can enhance host heat tolerance in specific groups, such as protecting bacteriocytes in aphids [105] or expressing thermotolerance genes in whiteflies [106], other have lost transcriptional regulators for temperature stimuli [107]. Such lineage-specific variations in endosymbiont or host interaction could explain how local environmental conditions (e.g. thermal variations across mesocosms) interact with host phylogeny. Additionally, unmeasured abiotic and biotic conditions, microbial isolation among mesocosms, or ecological drift may further contribute to these spatial patterns [108,109].

Given the observed phylogenetic signal, we investigated how climate and environmental factors influence on shorter time scales microbiota diversity and composition within families: two detritivorous (Isopodae, Julidae), one phytophagous (Cicadellidae) and one predatory (Lycosidae) arthropod families. We observed contrasted effects across arthropod families. In detritivores, the climatic treatment appeared as a prominent driver of host-associated microbiota diversity or composition, both directly or in interaction with other factors. However, its effect differed between the two detritivorous families. In millipedes, the microbiota diversity was directly and negatively affected by a warmer climate, and so was its composition which is in line with patterns usually observed in invertebrates [34]. Isopods responded differently, with climate having no clear direct effect both on diversity and composition, the latter being mostly driven by the abundance of isopods predators. This is in line with findings in daphnia, whose associated microbiota respond to predation stress [110].

For both detritivorous host families, we found that climate had interactive effects with resources-related variables. In Millipedes, it was with the host δ¹⁵N. Since δ¹⁵N enrichment could reflects accelerated decomposition and microbial nitrogen processing under heat, this isotopic marker may track changes in nitrogen-metabolizing bacteria such as *Microbacteriaceae* and *Devosia* [111]. However, data on millipede microbiota remain scarce in general and further studies are needed to test this hypothesis. In isopods, warming weakly increased microbiota diversity in habitats with high phytophagous richness and detritivore abundance, an effect disappearing under low richness and abundance conditions. This result contrasts with laboratory warming experiments, where isopod gut microbiota decreased [112]. Warmer conditions were also associated with slightly higher phytophagous richness and detritivore abundance. This may alter litter composition, resource availability, and thereby detritivore-associated microbiota [113], as shown in earthworms where diet shifts rapidly restructure microbial communities [114]. This suggests that climatic effects on microbiota diversity may be mediated indirectly through resource-driven changes in the host environments.

For the other trophic guilds, variables related to potential or effective resources were, in most cases, responsible for most of the explained variation in the microbiota diversity and composition. In Lycosidae hosts, who are predators, the microbiota diversity was positively influenced by detritivorous abundance while its composition was explained by phytophagous abundance, both groups being potential prey of spiders. Their microbiota composition was also explained by their own abundance, which may be explained by the fact that spider’s diet also relies on spiders, including conspecifics [115]. The observed lower microbiota diversity in Lycosidae individuals exhibiting higher δ¹⁵N values may hence reflect restricted diet and cannibalism.

The microbiota diversity of phytophagous hosts was overall negatively influenced by phytophagous richness and abundance. Higher abundance and aggregation of phytophagous hosts may increase competition for shared plant resources. This may impose greater competitive pressure on individual hosts and affect their microbiota. Alternatively, higher abundance in phytophagous hosts may increase microbial transmission among host individuals, leading to microbiota homogenization. The composition of microbiota associated with phytophagous hosts was mainly explained by ∂^13^C and ∂^15^N, which could reflect local resources and habitat. In leafhoppers, we observed a different trend though, with a small quadratic response of microbiota diversity to climate. Interestingly, climate was also the main drivers of bacterial phylogenetic dissimilarity for that host clade. A possible explanation is that warming reduces bacterial symbiont diversity and abundance, as found in the green stinkbug [36], though this effect may be buffered under thermal extremes. For example, the secondary endosymbiont *Sodalis* spp., found in cicadellids, include heat-tolerant symbionts (*Sodalis glossinidius*) that upregulate heat-shock genes around 30–37 °C in tse-tse flies [116]. A similar mechanisms could be at play, although this experimental validation is still required [117].

By studying arthropod microbiota under semi-natural conditions where we manipulated the biotic (presence or absence of an apex predator) and abiotic context (climatic treatment) and accounting for hosts phylogenetic and functional characteristics, we demonstrate that macro-scale evolutionary factors, such as host phylogeny and phylogenetically conserved traits, act as primary drivers of host microbiota at the scale of an entire metacommunity of arthropods, despite a fluctuating environmental context. On the other hand, we also identify finer-scale, ecological processes in host microbiota linked to the immediate biotic and abiotic context of the host. This duality of responses complicates the interpretation of global change impacts on host-microbiota associations. As climate change and trophic downgrading – corresponding to the two treatments applied here – impose increasing pressures on biodiversity, one could wonder at what level of climatic / biotic pressure these long-term associations would reach a tipping point or be disrupted. Unravelling these limits will be key to determining the threshold at which host microbiota associations translates into host physiological changes and altered functional dynamics for the entire ecosystem [118].

## Acknowledgements

The experiments were performed at the B2M facility, and we thank the support and resources provided by the B2M team, in particular Anne-Sophie Benoiston for her advice and help on bioinformatics analyses. We are also deeply grateful to Benjamin Linard for his advice on the phylogenetic placement analyses, as well as to the Genotoul bioinformatics platform Toulouse Occitanie (Bioinfo Genotoul, https://doi.org/10.15454/1.5572369328961167E12) for providing computing and storage resources. We also thank Luis San-José, Félix Pellerin and Laurane Winandy for their essential’s contribution to fieldwork. This work was supported by the French Laboratory of Excellence project ‘TULIP’ (grant nos ANR-10-LABX-41 and ANR-11-IDEX-0002-02), by an ‘Investissements d’avenir’ program from the Agence Nationale de la Recherche (grant no. ANR-24-INBS-0001 AnaEE-Services), and by the FRAIB Young Scientist 2022 program (project PREDABACT). This work is part of a project that has received funding from the European Research Council (ERC) under the European Union’s Horizon 2020 research and innovation program (grant agreement no. 817779 to J.Co.).

## Data availability statement

The datasets, metadata and scripts generated and analysed during the current study will be made available upon manuscript acceptance.

## Notes

### Competing Interest Statement

The authors have declared no competing interest.

