## Supplementary_material for "Disentangling the eco-evolutionary drivers of host-associated microbiota at the scale of an arthropod meta-community"

**Figure S1** Principal component analysis (PCA) and respective contribution and coordinates of each variable on the first two PCA axes for climatic variables within the Metatron.

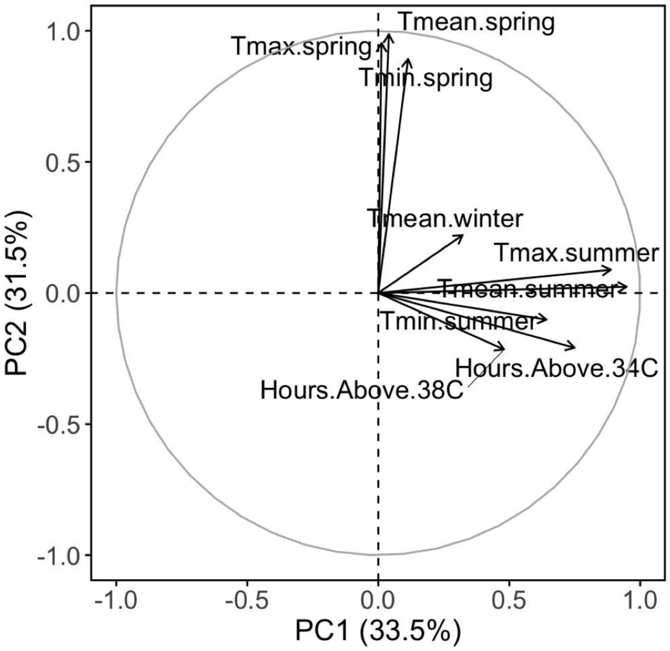

| | Contrib | Coord |
|---|---|---|
| **DIM1 (33.5%)** | | |
| Tmean.summer | 29.877 | +0.948 |
| Tmax.summer | 26.166 | +0.888 |
| Hours.Above.34C | 18.660 | +0.749 |
| Tmin.summer | 13.698 | +0.642 |
| Hours.Above.38C | 7.666 | +0.480 |
| Tmean.winter | 3.449 | +0.322 |
| Tmin.spring | 0.430 | +0.114 |
| Tmean.spring | 0.055 | +0.040 |
| Tmax.spring | 0.005 | +0.012 |
| **DIM2 (31.5%)** | | |
| Tmean.spring | 34.320 | +0.987 |
| Tmax.spring | 32.015 | +0.953 |
| Tmin.spring | 28.107 | +0.893 |
| Tmean.winter | 1.715 | +0.221 |
| Hours.Above.38C | 1.637 | -0.216 |
| Hours.Above.34C | 1.542 | -0.209 |
| Tmin.summer | 0.366 | -0.102 |
| Tmax.summer | 0.278 | +0.089 |
| Tmean.summer | 0.019 | +0.023 |

**Figure S2** A. Distribution of sample sizes by number of pooled individuals (1–2, 3, >3). B. Diversity metrics (Richness, Shannon index, and total reads) as a function of the number of pooled individuals. C. Sample dispersion based on Kantorovich-Rubinstein phylogenetic distances across pooling levels. D. Sample size and average distance to group centroid from panel C, estimated using the `permdisp` function (from the `vegan` R package).

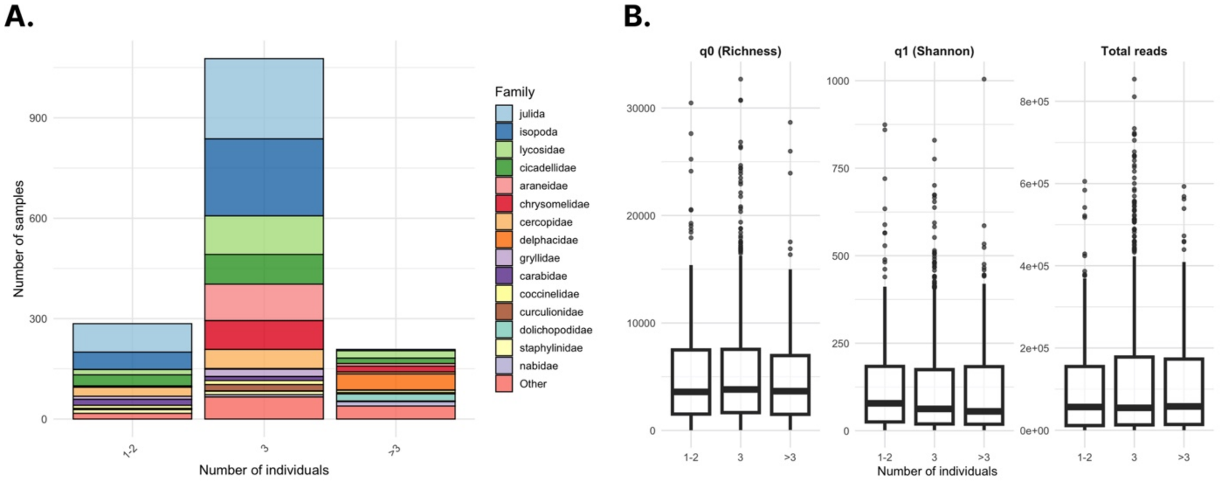

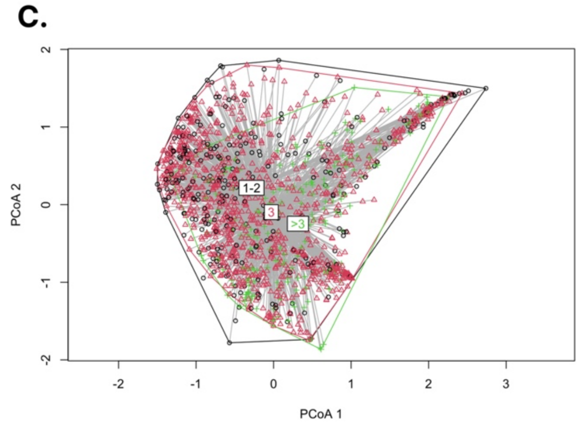

**D.**

| | 1-2 | 3 | <3 | NA |
|---|---|---|---|---|
| Sample size | 342 | 1077 | 151 | 5 |
| Average distance to median | 1.730 | 1.597 | 1.542 | |

**Figure S3** Rarefaction and sequencing completeness curves of each sample. Graphics respectively show the OTUs richness (D0), the exponential of Shannon index (D1), the inverse of the Simpson index (D2) and the Good's coverage index (coverage).

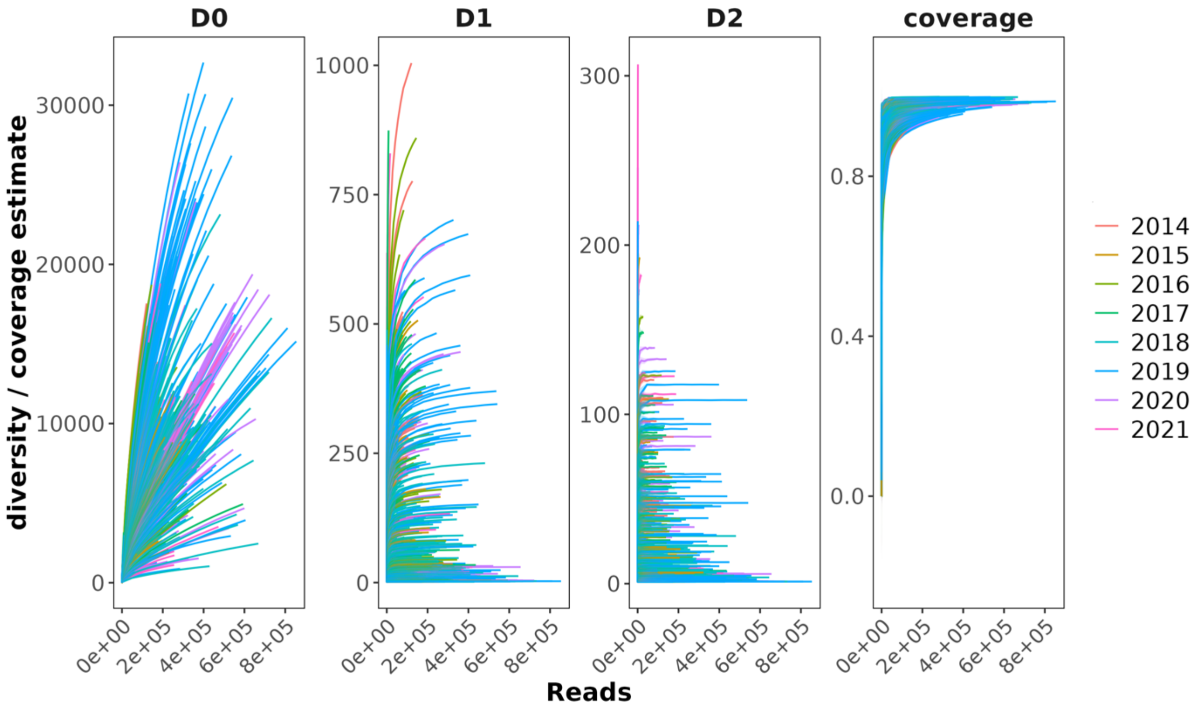

**Figure S4** Accumulated Likelihood Weight Ratio (LWR) values as a function of sequence index (p-query) for the first five placements. The x-axis represents sorted sequence indices (p-queries), while the y-axis shows the accumulated LWR at each p-query, showing the distribution of placement probabilities across the reference tree. On the left side of the plot, p-queries have nearly all their LWR in the most likely placement (LWR.1), indicating confident placement on a single branch. Most p-queries have their placement probability concentrated within the first three positions, suggesting minor ambiguity among a few branches but generally reliable placement. The thin yellow region on the right represents p-queries with higher placement uncertainty, as more of their probability is distributed in the "Remainder" category.

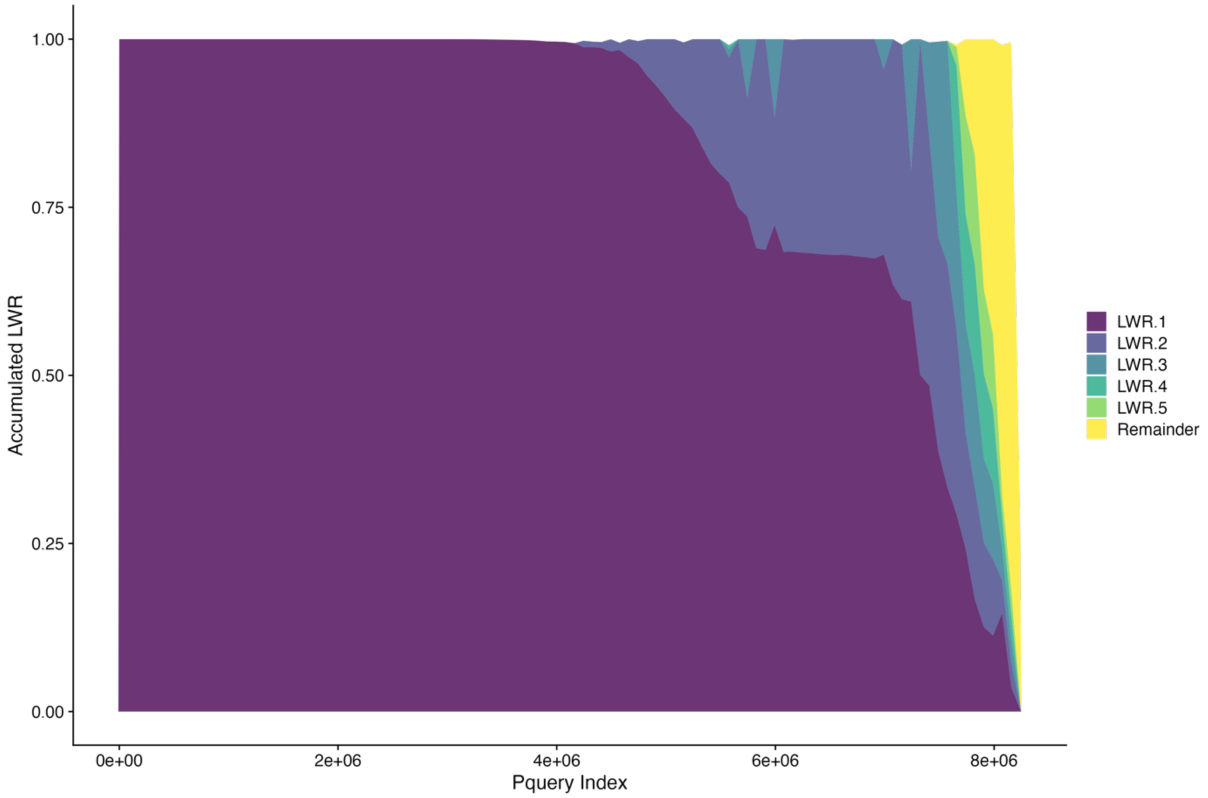

**Figure S5** Principal Coordinate Analysis of the isotopic signature dissimilarities between arthropod families based on Euclidean distances calculated from the mean $\delta^{15}N$ and $\delta^{13}C$ values for each family.

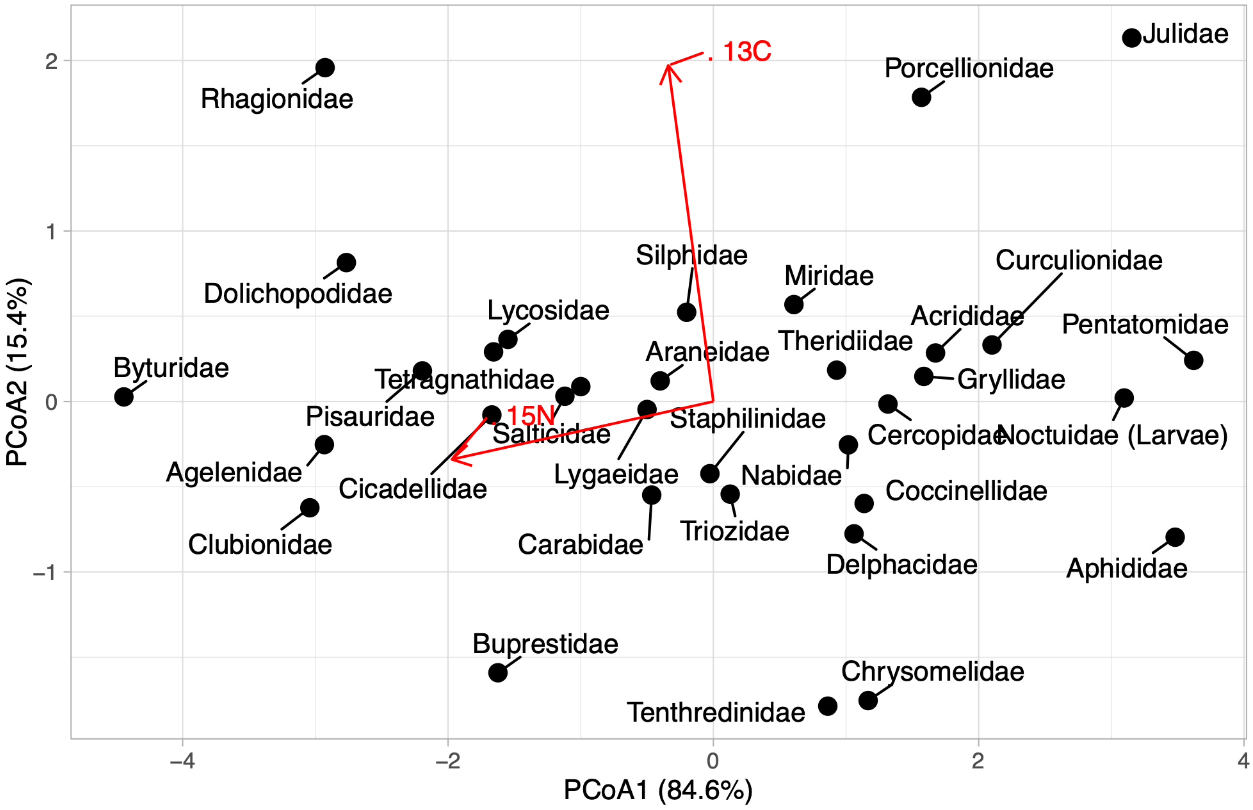

**Table S1** Sampled arthropods families by trophic group and their top 3 dominant microbiome genus.

| Family | Genus 1 | Genus 2 | Genus 3 |
|---|---|---|---|
| **Detritivorous (n=438)** | | | |
| Isopoda | Rickettsiella | Paracoccus | Wolbachia |
| Julidae | Candidatus Rhabdochlamydia | Phaeovibrio | Rickettsiella |
| **Phytophagous (n=359)** | | | |
| Acrididae | Rickettsiella | Leifsonia | Pseudomonas |
| Aphididae | Candidatus Regiella | Buchnera | Rickettsiella |
| Buprestidae | Vibrio | Serratia | Pseudoalteromonas |
| Byturidae | Wolbachia | Kluyvera | Pseudomonas |
| Cercopidae | Candidatus Sulcia | Candidatus Rhabdochlamydia | Rickettsiella |
| Chrysomelidae | Rickettsiella | Wolbachia | Rickettsia |
| Cicadellidae | Candidatus Sulcia | Pectobacterium | Rickettsiella |
| Curculionidae | Rickettsiella | Pantoea | Endosymbionts |
| Delphacidae | Rickettsia | Pantoea | Rickettsiella |
| Gryllidae | Lactococcus | Klebsiella | Enterococcus |
| Liviidae | Sodalis | Candidatus Carsonella | Nocardioides |
| Lygaeidae | Enterococcus | Rickettsia | Rickettsiella |
| Miridae | Rickettsia | Neisseria | Pseudomonas |
| Noctudiae (larva) | Serratia | Vibrio | Pseudoalteromonas |
| Pentatomidae | Tatumella | Agrococcus | Enterococcus |
| Tenthredinidae | Methylobacterium-Methylorubrum | Sphingomonas | Pseudomonas |
| **Predator (n=348)** | | | |
| Pisauridae | Rickettsiella | Rickettsia | Sphingomonas |
| Rhagionidae | Rickettsiella | Pseudoalteromonas | Sphingomonas |
| Salticidae | Parapusillimonas | Diplorickettsia | Rickettsiella |
| Silphidae | Tyzzerella | Knoellia | Enterococcus |
| Staphylinidae | Pseudomonas | Amycolatopsis | Rickettsiella |
| Tetragnathidae | Enterococcus | Rickettsia | Buchnera |
| Theridiidae | Wolbachia | Candidatus Cardinium | Spingomonas |
| Thomisidae | Candidatus Fritschea | Staphylococcus | Rickettsiella |
| Agelenidae | Rickettsia | Sphingomonas | Afipia |
| Lycosidae | Rickettsiella | Diplorickettsia | Candidatus Fritschea |
| Araneidae | Rickettsiella | Diplorickettsia | Candidatus Rhabdochlamydia |
| Clubionidae | Streptococcus | Candidatus Sulcia | Spingomonas |
| Coccinellidae | Rickettsiella | Wolbachia | Buchnera |
| Dolichopodidae | Rickettsiella | Aeromonas | Spiroplasma |
| Nabidae | Rickettsiella | Buchnera | Enterococcus |

**Table S2** Summary of models explaining the presence of Wolbachia and Rickettsia. Fixed effects include climatic treatment (Climate t-1), arthropods richness, lizard density and host phylogenetic position (PCoA1), mobility (evasive, intermediate or sedentary) and trophic guild (detritivores, phytophagous, predators). Mesocosms, years and families were included as random intercepts. Parameters not shown in the table were excluded from the models based on the AIC selection procedure.

| Parameters | Estimate | SE | z-value | RI | p-value |
|---|---|---|---|---|---|
| **Wolbachia** $R^2_m = 0.049$; $R^2_c = 0.221$ | | | | | |
| Intercept | 2.645 | 0.897 | 2.947 | 1 | 0.003** |
| Climate (t-1) | -0.161 | 0.113 | 1.422 | 0.92 | 0.155 |
| Host distance | 0.309 | 0.152 | 2.033 | 0.46 | 0.042* |
| Mobility: intermediate | -1.462 | 1.065 | 1.371 | 0.18 | 0.170 |
| Mobility: sedentary | -1.579 | 1.119 | 1.409 | 0.26 | 0.159 |
| Arthropods richness | 0,109 | 0.168 | 0.651 | 0.26 | 0.515 |
| Lizard density | 0.032 | 0.116 | 0.272 | 0.16 | 0.785 |
| ***Random effects*** | | | | | |
| Family | 0.219 | 0.468 | | | |
| Mesocosm ID | 0.014 | 0.118 | | | |
| Year | 0.496 | 0.705 | | | |
| **Rickettsia** $R^2_m = 0.038$; $R^2_c = 0.120$ | | | | | |
| Intercept | 1.048 | 0.183 | 5.704 | 1 | <0.001** |
| Arthropods richness | 0.340 | 0.150 | 2.258 | 1 | 0.024* |
| Lizard density | -0.138 | 0.088 | 1.563 | 0.63 | 0.118 |
| Climate (t-1) | -0.186 | 0.083 | 2.231 | 1 | 0.026* |
| Host distance | 0.079 | 0.155 | 0.512 | 0.19 | 0.609 |
| ***Random effects*** | | | | | |
| Family | 0.236 | 0.486 | | | |
| Mesocosm ID | 0.072 | 0.269 | | | |

**Table S3** Summary of models explaining bacterial diversity across the ubiquitous families: Julidae (N=201), Isopoda (N= 211), Cicadellidae (N=84) and Lycosidae (N=109). Fixed effects include climatic treatment (Climate t-1), arthropods richness and abundance per trophic group, plants species richness, lizard density, lizard biomass per density. Mesocosms, years and families were included as random intercepts. Interactions and parameters not shown in the table were excluded from the models based on the AIC selection procedure.

| Parameters | Estimate | SE | z-value | RI | p-value |
|---|---|---|---|---|---|
| **Julidae** $R^2_m = R^2_c = 0.129$ | | | | | |
| Intercept | 121.863 | 12.144 | 9.977 | 1 | <0.001*** |
| $\partial^{13}C$ | 16.370 | 10.320 | 1.577 | 0.81 | 0.115 |
| $\partial^{15}N$ | 26.198 | 12.350 | 2.109 | 1 | 0.035* |
| Climate (t-1) | -40.327 | 11.804 | 3.396 | 1 | <0.001*** |
| Climate (t-1) * $\partial^{15}N$ | -36.708 | 13.276 | 2.748 | 1 | 0.006** |
| Plant richness | 11.432 | 9.859 | 1.152 | 0.24 | 0.249 |
| Phytophagous richness | -9.403 | 8.900 | 1.050 | 0.09 | 0.294 |
| Climate (t-1) * $\partial^{13}C$ | -12.798 | 11.825 | 1.076 | 0.20 | 0.282 |
| I (Climate $(t-1)^2$) | -7.138 | 7.165 | 0.990 | 0.14 | 0.322 |
| Lizard density | 5.705 | 9.230 | 0.614 | 0.06 | 0.539 |
| Predator abundance | -5.385 | 10.857 | 0.493 | 0.06 | 0.622 |
| Detritivorous abundance | -3.522 | 9.209 | 0.380 | 0.06 | 0.704 |
| **Isopodae** $R^2_m = R^2_c = 0.165$ | | | | | |
| Intercept | 99.548 | 9.384 | 10.547 | 1 | <0.001*** |
| Phytophagous richness | 14.348 | 10.561 | 1.351 | 1 | 0.176 |
| Predator richness | 15.252 | 9.341 | 1.623 | 0.70 | 0.105 |
| Climate (PC1) | 13.935 | 9.358 | 1.480 | 1 | 0.139 |
| Detritivorous abundance | -4.002 | 10.040 | 0.396 | 1 | 0.692 |
| Phytophagous abundance | -16.471 | 10.471 | 1.564 | 0.69 | 0.118 |
| Climate (t-1) * Phytophagous richness | 27.214 | 9.604 | 2.817 | 1 | 0.005** |
| Climate (t-1) * Detritivorous abundance | 44.130 | 12.528 | 3.502 | 1 | <0.001*** |
| Lizard biomass/density | -4.673 | 8.942 | 0.519 | 0.23 | 0.603 |
| Climate (t-1) * Lizard biomass/density | -16.047 | 10.274 | 1.553 | 0.16 | 0.120 |
| $\partial^{15}N$ | 7.910 | 10.506 | 0.748 | 0.15 | 0.454 |
| Climate (t-1) * Phytophagous abundance | -4.545 | 8.820 | 0.512 | 0.07 | 0.608 |
| **Cicadellidae** $R^2_m = 0.231$, $R^2_c = 0.275$ | | | | | |
| Intercept | 82.041 | 25.350 | 3.204 | 1 | 0.001** |
| I (Climate $(t-1)^2$) | 20.264 | 10.221 | 1.953 | 0.84 | 0.051. |
| Detritivorous abundance | -16.936 | 11.955 | 1.395 | 0.34 | 0.163 |
| $\partial^{13}C$ | 29.585 | 24.149 | 1.206 | 0.18 | 0.228 |
| $\partial^{15}N$ | 20.600 | 18.298 | 1.109 | 0.15 | 0.267 |
| Climate (t-1) | 11.671 | 13.010 | 0.883 | 0.08 | 0.377 |
| Lizard density | -12.900 | 13.745 | 0.924 | 0.08 | 0.355 |
| Phytophagous richness | -8.616 | 12.284 | 0.691 | 0.07 | 0.490 |
| ***Random effect*** | | | | | |
| Mesocosm ID | 5.7% | | | | |
| **Lycosidae** $R^2_m = 0.136$; $R^2_c = 0.166$ | | | | | |
| Intercept | 114.815 | 17.096 | 6.644 | 1 | <0.001*** |
| $\partial^{15}N$ | -35.957 | 17.980 | 1.977 | 0.92 | 0.048* |
| Detritivorous abundance | 18.149 | 11.461 | 1.565 | 0.72 | 0.117 |
| Phytophagous richness | -11.759 | 11.749 | 0.989 | 0.19 | 0.323 |
| Predator abundance | -9.480 | 11.829 | 0.792 | 0.09 | 0.428 |
| I (Climate $(t-1)^2$) | 6.387 | 9.398 | 0.672 | 0.09 | 0.502 |
| Predator richness | -6.343 | 11.850 | 0.529 | 0.08 | 0.597 |
| Lizard biomass/density | 5.534 | 11.069 | 0.494 | 0.08 | 0.621 |
| ***Random effect*** | | | | | |
| Year | 3.4% | | | | |

### Appendix A: Details of PCR mix components and thermocycling conditions.

PCR reactions were conducted in triplicate for each sample in 20 µl containing 3 µl of 1/10 diluted DNA extract, 0.25 µM of each primer, 1 U of AmpliTaq Gold DNA Polymerase (Applied Biosystems, Foster City, CA, USA), 2.5 mM of MgCl2, 1x of Taq Buffer, 0.2 mM of each dNTP and 4 ng of bovine serum albumin (Promega Corporation, Madison, USA). PCR thermocycling conditions were as follows: 5 min of initial denaturation at 95°C, followed by 35 cycles of denaturation (95°C for 30 s), annealing (57°C for 30 s), elongation (72°C for 30 s) and a final elongation (72°C for 7 min).

### Appendix B: Identifying the drivers of arthropod microbiota taxonomic diversity and composition at the trophic guild level.

#### Methods

Detritivores were excluded, as analyses had already been conducted separately for the two sampled families. Trophic guilds integrate ecological differences beyond diet, including lifestyle, microhabitat, and exposure to distinct microbial sources, which may influence microbiota composition. Previous work [1,2] supports the relevance of guild-level effects at the whole-organism scale. Detritivores were excluded, as analyses had already been conducted separately for the two sampled families.

Trophic guild-level analyses were conducted using the same model-averaging approach, fixed and random effect structure, climate interactions, and model selection procedure as described in the main manuscript for the family-level analyses. However, to account for the effects of host phylogeny and mobility at this broader scale, we included host phylogenetic distance (represented by the first axis of the PCoA on host phylogenetic distance, explaining 47.16 % of the variance) and mobility potential as fixed effects, and arthropod family as a random effect.

#### Results

When extending the analysis to the trophic guild level, we found no significant differences in microbiota alpha diversity between trophic guilds or trophic positions (Fig. S6), and host family and phylogeny explained little to no variance (Table S4). Overall, the explanatory variables accounted for less variance in microbiota diversity compared to the family-level models, as indicated by lower variable relative importance and lower $R^2$ values. However, the microbiota diversity of predatory arthropods tended to be positively associated with detritivores abundance, a trend also observed in the Lycosidae family (Table S3). In phytophagous arthropods, phytophagous richness and abundance tended to be negatively associated with phytophagous diversity. We found that the effect of predator abundance depended on the climate with a positive interaction. Mobility capacity was retained in neither the phytophagous nor the predator models (Table S4).

At the trophic guild levels, GDM models still explained a low fraction of the total phylogenetic deviance (Fig. S7). For the microbiota of predatory arthropods, the deviance was prominently explained by predator abundance and year. Finally, the deviance in phytophagous microbiota phylogenetic dissimilarities were best explained by the host $\delta^{13}C$ and then $\delta^{15}N$ content, as well by the lizard biomass-density ratio.

**Figure S6** Microbiota diversity according to trophic position defined by (A) trophic guild and (B) isotopic measures ($\delta^{13}C$ representing carbon source, and $\delta^{15}N$ indicating the trophic position of the sample).

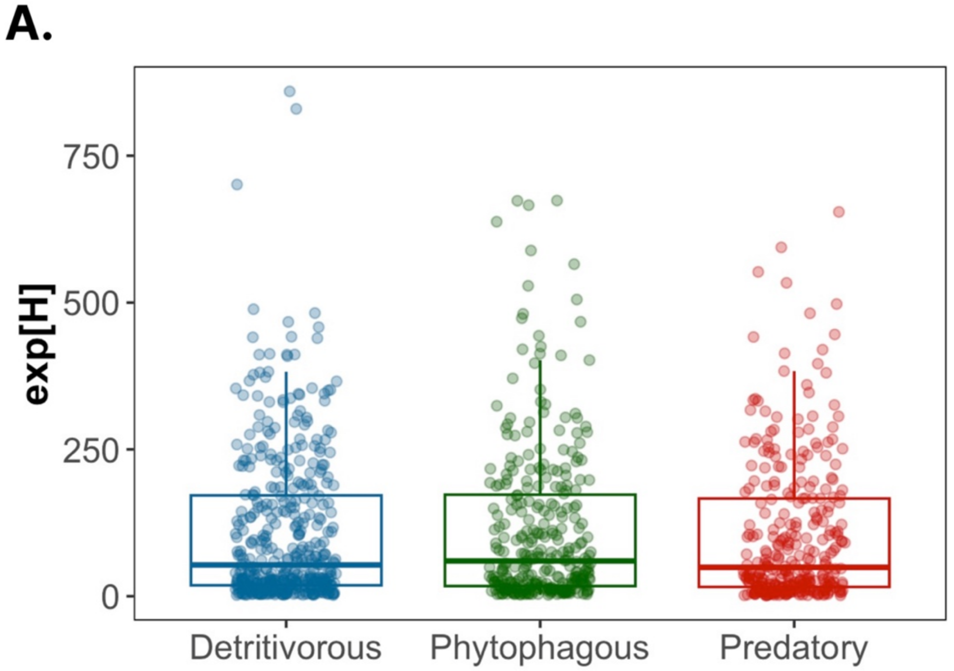

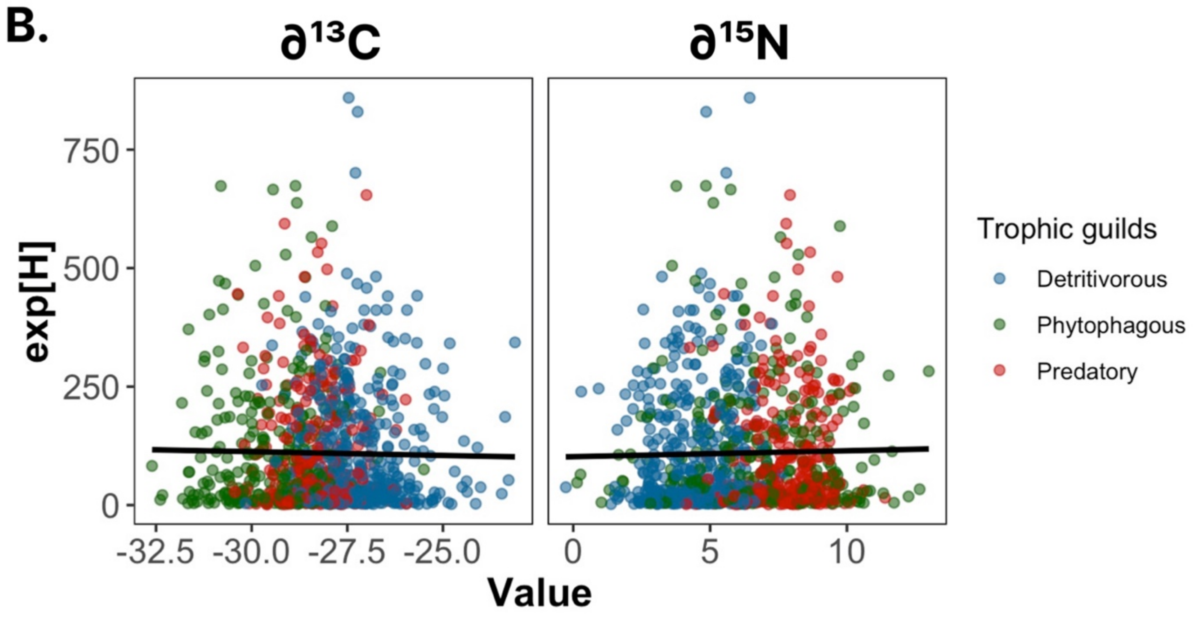

**Figure S7** Generalized dissimilarity models (GDMs) splines for arthropod's trophic guild (Predators and Phytophagous). We used Climate (t-1), arthropods diversity and abundance and isotopic measurement, as well as lizard diversity and plant richness. The shape of the spline shows the rate of community composition changes (y-axis) as the predictor variables (x-axis) increases. Predictors variables are scaled. Numbers indicate the percentage of deviance explained.

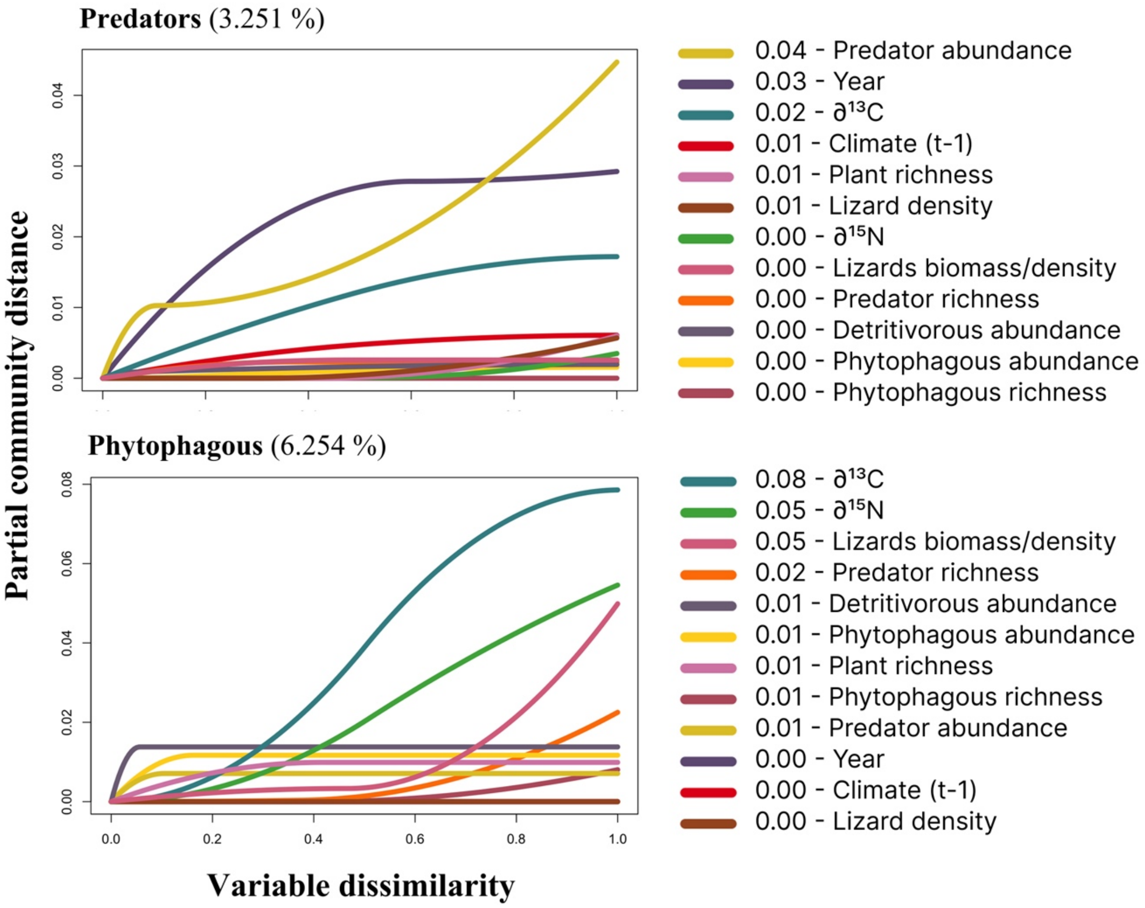

**Table S4** Summary of models explaining bacterial diversity across trophic groups. Fixed effects included climatic treatment (Climate (t-1)) and its quadratic term (I(Climate $(t-1)^2$)), arthropod richness and abundance per trophic group, plant species richness, lizard density, and lizard biomass per density. Mesocosms, years, and families were included as random intercepts. Host relatedness was included as a covariate only in the phytophagous model, as it was redundant with the family variable in the detritivorous model (which had only two levels) and highly collinear with mobility in the predator model. Interactions and parameters not shown in the table were excluded from the models based on AIC model selection. Only random effects that contributed to variance were retained for model averaging.

| Parameters | Estimate | SE | z-value | RI | p-value |
|---|---|---|---|---|---|
| **Predators** $R^2_m = 0.065$; $R^2_c = 0.076$ | | | | | |
| Intercept | 103.345 | 9.839 | 10.464 | 1 | <0.001*** |
| Phytophagous richness | -11.705 | 7.235 | 1.611 | 0.57 | 0.107 |
| Detritivorous abundance | 14.973 | 7.474 | 1.995 | 0.91 | 0.046* |
| Climate (t-1) | -9.042 | 7.562 | 1.191 | 0.21 | 0.234 |
| Lizard biomass | 9.224 | 7.756 | 1.185 | 0.30 | 0.236 |
| I (Climate $(t-1)^2$) | 6.524 | 5.584 | 1.163 | 0.25 | 0.245 |
| Host distance | -7.070 | 6.861 | 1.026 | 0.19 | 0.305 |
| Plant richness | 6.730 | 7.646 | 0.877 | 0.07 | 0.381 |
| Predator richness | 5.712 | 7.480 | 0.761 | 0.06 | 0.447 |
| Lizard density | -9.804 | 8.584 | 1.138 | 0.06 | 0.255 |
| $\partial^{13}C$ | -3.273 | 7.006 | 0.465 | 0.04 | 0.642 |
| Phytophagous abundance | -3.612 | 7.175 | 0.501 | 0.02 | 0.616 |
| $\partial^{15}N$ | 2.498 | 6.922 | 0.359 | 0.04 | 0.719 |
| ***Random effects*** | | | | | |
| Year | 1.17% | | | | |
| **Phytophagous** $R^2_m = R^2_c = 0.058$ | | | | | |
| Intercept | 112.643 | 22.899 | 4.903 | 1 | <0.001*** |
| Phytophagous richness | -17.85 | 9.415 | 1.889 | 0.81 | 0.059. |
| Phytophagous abundance | -17.845 | 9.195 | 1.859 | 0.86 | 0.063. |
| Lizard density | -10.981 | 7.902 | 1.384 | 0.40 | 0.166 |
| Detritivorous abundance | -12.247 | 8.355 | 1.460 | 0.36 | 0.144 |
| Climate (t-1) | 2.283 | 8.496 | 0.268 | 0.11 | 0.789 |
| Predator abundance | 17.645 | 16.685 | 1.055 | 0.14 | 0.291 |
| Climate (t-1) * Predator abundance | 22.579 | 9.497 | 2.368 | 0.11 | 0.018 |
| Predator richness | 6.231 | 8.309 | 0.747 | 0.11 | 0.445 |
| Plant richness | -5.852 | 7.869 | 0.741 | 0.11 | 0.459 |
| Host distance | 0.181 | 0.372 | 0.483 | 0.04 | 0.629 |
| $\partial^{15}N$ | 2.566 | 7.775 | 0.329 | 0.03 | 0.742 |
| $\partial^{13}C$ | -2.553 | 7.795 | 0.326 | 0.03 | 0.744 |
